# BatCRISPRi: Bacillus titratable CRISPRi for dynamic control in *Bacillus subtilis*

**DOI:** 10.1101/2023.11.01.565046

**Authors:** Andrés F. Flórez, Sebastian Castillo-Hair, Luis Gutierrez-Lopez, Daniel Eaton, Johan Paulsson, Ethan C. Garner

**Affiliations:** Department of Molecular and Cellular Biology, Harvard University, Cambridge, MA, USA; Department of Electrical and Computer Engineering, University of Washington, Seattle, WA; eScience Institute, University of Washington, Seattle, Washington 98195, United States; Department of Systems Biology, Harvard Medical School, Boston, MA, USA

## Abstract

The discovery of new genes regulating essential biological processes has become increasingly important, and CRISPRi has emerged as a powerful tool for achieving this goal. This method has been used in many model organisms to decrease the expression of specific genes and assess their impact on phenotype. Pooled CRISPRi libraries in bacteria have been particularly useful in discovering new regulators of growth, division, and other biological processes. However, these libraries rely on the induction of dCas9 via an inducible promoter, which can be problematic due to promoter leakiness. This is a widespread phenomenon of any inducible promoter that can result in the unwanted downregulation of genes and the emergence of genetic suppressors when essential genes are knocked down. To overcome this issue, we have developed a novel strategy that eliminates dCas9 leakiness and enables reversible knockdown control using the rapamycin-dependent degron system in *Bacillus subtilis*. This degron system causes rapid degradation of dCas9, resulting in an almost instant reset of the system. Our results demonstrate that it is possible to achieve zero CRISPRi activity in the uninduced state and full activity in the induced state. This improved CRISPRi system will enable researchers to investigate phenotypic changes more effectively while reducing the undesirable effects of leaky expression and noise in their phenotypic data. Moreover, a rapid degradation system could serve as a tool for dynamic perturbation before compensation mechanisms or stress responses kick in. Finally, this approach can be adapted to other organisms and other promoter-inducible systems, potentially opening up strategies for tighter control of gene expression.

## Introduction

Clustered Regularly Interspaced Short Palindromic Repeats (CRISPR) has revolutionized the field of molecular biology and genetics by enabling precise and efficient genome editing in various organisms^1,2^. The CRISPR system comprises a Cas9 nuclease and a guide RNA that directs the Cas9 to the specific target sequence, which induces double-strand breaks (DSBs) in the DNA^3^. However, the CRISPR system has also been adapted for transcriptional regulation. In CRISPR interference (CRISPRi), the Cas9 nuclease is deactivated, and the guide RNA (gRNA) directs it to the promoter region of the target gene, leading to transcriptional repression^4^. The versatility and simplicity of this technique have made it a popular tool for gene expression studies^5,6^, metabolic engineering^7,8^, and therapeutic applications^9^.

CRISPRi offers several advantages over traditional gene silencing techniques, such as RNA interference (RNAi) and overexpression^10^. These advantages include specificity, efficiency, and reversibility. However, CRISPRi systems expressing dCas9 from inducible promoters have a fundamental problem: leakiness, resulting from an inherent limitation of inducible promoters^11^. Even a few molecules of dCas9 can cause transcriptional repression in the absence of the inducer due to the high binding efficiency of dCas9^12^. For *Bacillus subtilis*, this resulted in 3-fold repression without dCas9 induction^5^. Several approaches have been used to address this problem, mainly at the post-transcriptional level. These approaches include promoter optimization^13^, mismatch gRNAs^14^, synthetic amino acids to control translation^15^, optogenetic CRISPRi^16^, and split dCas9^17^. However, none of these approaches has completely eliminated leakiness, and their engineering is complex or the presence of phototoxicity in the case of the optogenetic system. For example, adding other genetic components may be required to use synthetic amino acids, and a specific gRNA may be needed for each level of repression, making the approaches cumbersome to implement^15^. Additionally, these approaches do not provide full reversibility which is important because it allows the study of the effect of gene expression on a particular phenotype in a temporal manner. Therefore, a more robust and simple-to-implement solution is needed.

A potential strategy to solve this issue is the use of degrons. A degron is a small peptide sequence that can be recognized by the cellular degradation machinery, leading to the rapid degradation of the tagged protein^18,19.^ There is an existent system already developed in *Bacillus subtilis* using the SsrA tag from *E*.*coli*^20^. By fusing a degron sequence to dCas9, it is possible to eliminate its leaky expression. Some attempts have been made in this direction^21^. The reversible nature of this system is particularly important, as it enables the restoration of gene expression upon removal of the degron inducer molecule. In this paper, we use a rapamycin-inducible degron system (here named rapamycin system throughout the study), reengineered from an existent system in *Escherichia coli* ^22^, to eliminate CRISPRi leakiness completely and allow for a rapid reversible deactivation. The system allowed for full repression with low noise. We believe this system will be important for having a tight, reliable, reversible CRISPRi system in *Bacitllus subtilis* that could be extended to other organisms.

## Results

### Fine-tuning CRISPRi for leakiness elimination

We began by adapting the CRISPRi design previously used in *B. subtilis*^5^. In this design, dCas9 is driven by the xylose inducible promoter (Pxyl), while the gRNA is constitutively expressed using the constitutive Pveg promoter (in short, veg). We used mApple fluorescence protein instead of RFP (to improve signal brightness) as a gene readout and tagged dCas9 with mNeonGreen to monitor its level. Essentially, the complex dCas9/gRNA forms close to the 5’ region of the mApple gene, preventing transcription as dCas9 is induced. A schematic of these designs is presented in Fig. 1a. Consistent with previous reports of this system, this initial design (without SsrA tag) resulted in high levels of basal knockdown in the absence of dCas9 induction compared to the no gRNA control (Fig. 1b). The robust knockdown before induction and the considerable variation in fluorescence distribution observed in the flow cytometry data (Fig. 1b) are attributed to the leakiness of dCas9.

**Figure 1.**
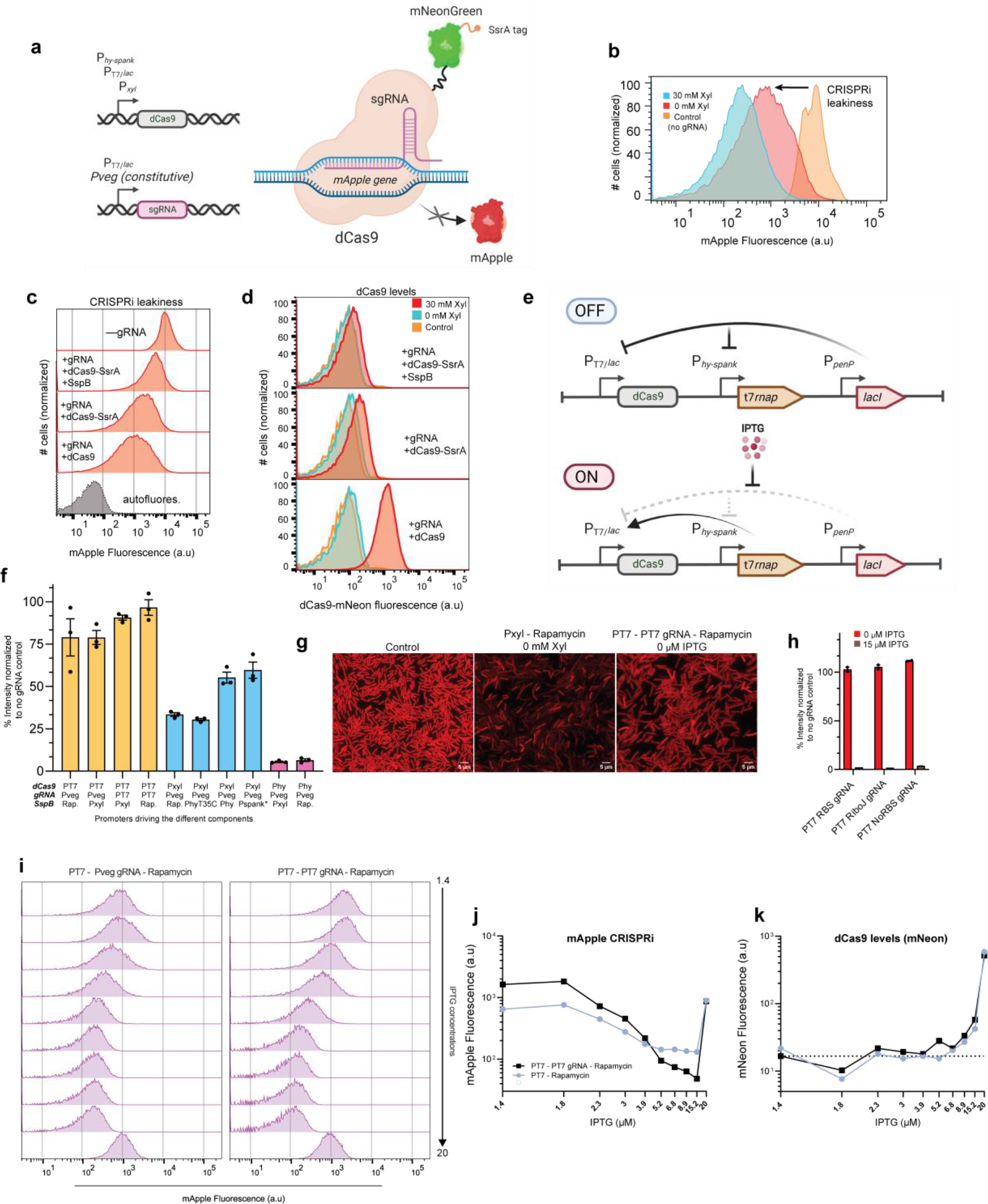
Fine-tuning CRISPRi for leakiness elimination. **a**, Schematic diagram of the CRISPRi system. dCas9 expression is driven by three different promoters (described in detail in the text). gRNA is driven by either a constitutive (veg) or inducible (T7) promoter. dCas9 blocks the expression of mApple and is tagged with mNeonGreen for intensity measurements, also displaying the SsrA tag. (Figure generated with BioRender). **b**, Flow cytometry measurements of Pxyl-dCas9 in the presence or absence of Xylose using a strain without gRNA as a control to assess leakiness; a representative experiment is shown. **c**, Leakiness measurement in different strains when adding an SsrA tag and SspB compared to the no gRNA control. mApple fluorescence recovery indicates a reduction in leakiness throughout the study. **d**, Fluorescence measurements of dCas9 levels measured simultaneously in **c**. Higher mNeon fluorescence indicates higher dCas9 levels. **e**, Schematic diagram of the T7 promoter and the new design including dCas9 and the switching mechanism upon addition of IPTG. dCas9 was replaced by gRNA when appropriate (Figure generated with BioRender). **f**, Quantification of leakiness from flow cytometry measurements with all the different genetic constructs, mixing a dCas9, gRNA, and SspB construct with the promoters listed in **a**. A 100% value means no leakiness. The first promoter is driving dCas9 and the second promoter drives SspB unless T7 gRNA is indicated, which means T7 is driving the gRNA expression (n= 3 biological replicates). **g**, Live cell snapshots on agar pads of the least leaky construct vs. control and Pxyl constructs. T7-T7 gRNA-Rapamycin exhibits high fluorescence homogeneity, indicating less leakiness compared to the Pxyl construct. **h**, Quantification via flow cytometry of leakiness of the T7 – T7 gRNA - Rapamycin construct while keeping the RBS, deleting it or replacing it by RiboJ (n= 3 biological replicates). **i**. IPTG titration of T7 – Rapamycin and T7-T7 gRNA-Rapamycin constructs looking at cell distributions in flow cytometry. **j**, Quantification of median mApple fluorescence from **i** and the corresponding dCas9 levels in **k**

Existing strategies for controlling leakiness in promoter constructs typically involve designing tighter promoters. While this approach can reduce leakiness, it may also limit the dynamic range of the promoter since tighter promoters are often weaker. We attempted an alternative strategy to achieve leakiness reduction by using protein degradation (Fig. 1a). The advantage of this strategy is that basal leaky degradation could eliminate leaky expressed dCas9 molecules while also allowing for reversible CRISPRi, where the gene can be reactivated after dCas9 is degraded. To achieve this, we added an SsrA (LGG) tag (abbreviated as SsrA*) to dCas9 (Fig. 1a) and monitored the levels of basal CRISPRi. The SsrA* tag binds to the SspB adaptor and promotes fast degradation of SsrA* protein fusions in *B. subtilis*^20^. Because there is leaky degradation in the absence of SspB, we hypothesized that adding SspB under a leaky inducible promoter might improve CRISPRi leakiness further by reducing free unwanted dCas9 resulting from leaky promoter expression. Adding only the SsrA tag slightly improved leakiness, while adding the Pspank*-SspB-driven system reduced leakiness even further compared to the no gRNA control (Fig. 1c). These results indicate that leaky degradation could counteract CRISPRi leakiness.

As depicted in Fig. 1d, the use of the SsrA* tag on dCas9 and the introduction of the inducible SspB construct led to a significant reduction in dCas9 leakiness. Nonetheless, we observed almost no dCas9 induction upon addition of the ligand, suggesting that the strength of the Pxyl promoter alone was insufficient to overcome the basal degradation rate (Fig. 1d). This made it challenging to silence highly expressed genes that required higher dCas9 occupancy. Since we encountered either persistent CRISPRi leakiness and lack of dCas9 inducibility, we explored alternative strategies. In *E. coli*, LacI-T7 promoters are frequently utilized for high protein production and are often less leaky than other promoters^11^. Recently, a LacI-T7 promoter was engineered in *B. subtilis* that exhibited a 2.7x higher expression with IPTG-relative to maximal induction of Phy-spank, but 31x lower expression without IPTG, exhibiting a significant improvement in leakiness^23^. Therefore, we reasoned that using a tighter promoter with a higher dynamic range could help address the issues we encountered with the Pxyl promoter. Hence, we used the LacI-T7 promoter to drive dCas9 and included Phy-spank, in addition to Pxyl, for comparison purposes. For a schematic representation of the design see Fig. 1e. In brief, LacI is expressed under a constitutive promoter (PpenP), while the T7 RNA polymerase is under the control of the Phy-spank promoter. Upon IPTG induction, T7 RNA polymerase is expressed, which then leads to the expression of dCas9.

We conducted experiments to assess the effect of different constructs on the leakiness of the CRISPRi system, using mApple signal as a readout. The main goal of our experiment was to identify which promoter from SspB was leaky enough to eliminate the dCas9 leakiness and whether inducing or not inducing the gRNA influenced dCas9 leakiness. Additionally, we aimed to identify a promoter that was strong enough to overcome the degradation rate and provide an appropriate dynamic range of dCas9 expression. For that purpose, we used three different constructs: (1) promoters driving dCas9 (PT7, Pxyl, or Phy-spank), (2) whether gRNA is driven by T7 (inducible) or Pveg (constitutive), and (3) which promoter was used to drive SspB expression (Pxyl, Rapamycin-inducible, Phy-spank T35C (discovered in this study and explained in the degron section), Phy-spank, Pspank*). We then combined a dCas9, gRNA, and SspB construct, ensuring that the inducers were compatible when combining the different constructs. To measure leakiness, we normalized the fluorescence intensity relative to the control (no gRNA), such that 100% intensity indicates no leakiness in the absence of the inducer.

We then assayed the leakiness of each construct by flow cytometry measurements. (Fig. 1f). The leakiest construct was Phy-spank as expected, and the promoters driving SspB weren’t leaky enough to significantly counteract the leakiness of Phy-spank. The Pxyl promoter retains some leakiness even with the degron system present. This implies that the promoter leakiness is strong even when suppressing it with degradation. Using very leaky promoters like Phy-spank for driving SspB expression only partially reduces Pxyl leakiness. However, stronger SspB leakiness with either Phy-spank or Pspank* reduced the overall CRISPRi leakiness. Using the T7 promoter strongly reduced the leakiness since this promoter is tighter than Phy-spank and Pxyl. When we add the gRNA as an inducible element (using the PT7 promoter), we surprisingly observed a reduction in leakiness to zero in the presence of the rapamycin-inducible system (which will be discussed in the next section). This result was similarly observed during live microscopy, where a heterogeneous mix of bright and dim cells was evident under Pxyl conditions, contrasting with the uniform fluorescence observed in the T7-T7 gRNA-Rapamycin construct (Fig. 1g). Hence, the combination of inducing gRNA only when needed, along with a very tight dCas9 promoter, outperforms the use of a tight dCas9 promoter alone. Based on these results, we conclude that by using a strong and tight promoter, in combination with inducible gRNA (also tightly regulated) and moderate degradation, we can achieve a perfect system with zero leakiness. This is achieved on the PT7-dCas9, PT7-gRNA and Rapamycin-SspB system.

We then tested whether sequences around the gRNA were crucial for its proper expression under the T7 promoter. We retained the strong RBS (MF001) originally present in the system, which should not be necessary for a gRNA. However, a study has shown that the absence of an RBS can affect RNA stability^24^ and thus may influence expression of our gRNA. We compared this to using RiboJ, a ribozyme that self-cleaves, thereby removing upstream sequences and eliminating promoter-associated elements. We also compared it to gRNA without any additional sequences. When assessing leakiness in comparison to full CRISPRi knockdown (15 μM IPTG), we did not observe any substantial differences. All constructs exhibited similar behavior, as illustrated in Fig. 1h. This finding implies that to reduce leakiness, no specific sequence is necessary upstream of the gRNA, so we opted to use the RBS version for the remaining experiments.

We next assessed the dose-response CRISPRi effect. To investigate this, we conducted a titration curve experiment comparing two systems: T7-dCas9, T7-gRNA, and Rapamycin-SspB versus the same system with Pveg-gRNA. The goal was to observe any differences when using an inducible gRNA. Both systems exhibited similar behavior; however, the system with Pveg-gRNA reached a state of CRISPRi saturation, indicating that further knockdown was not achievable (Fig. 1i,j). This observation is intriguing, as the *B. subtilis* CRISPRi study demonstrated that Pxyl was sufficient to induce full knockdown^5^. It is possible that due to the system’s leakiness and the potentially low expression of the readout gene, this level of knockdown was attainable. In our system, mApple is highly expressed, necessitating higher levels of dCas9 and gRNA to achieve complete knockdown. While knowing the specific stoichiometry of dCas9-gRNA required for full knockdown at very high expression levels is beyond the scope of this paper, it is noteworthy that at the highest inducer levels the CRISPRi effect was abolished (Fig1.j), regardless of gRNA levels. Upon examination of dCas9 levels, we observed exceptionally high expression at an IPTG dose of 20 μM (Fig. 1k). It is conceivable that at these elevated levels, some form of phase separation may hinder dCas9 from binding to its target due to the formation of non-functional protein aggregates. However, further experiments are necessary to test this hypothesis, as other systems typically do not employ such high dCas9 levels. The possibility of phase separation-induced effects on dCas9 functionality warrants further investigation.

### Building a rapid degradation system through dimerization in *Bacillus subtilis*

As stated before, we used a rapamycin-inducible system to degrade dCas9, eliminate leakiness, and make the system reversible by degrading dCas9 and re-activating the gene after the knockdown. The rapamycin system was developed in *Bacillus subtilis* in this study. The system, which was first described in *E. coli*^22^, involved using the rapamycin-binding domain from mTOR called FRB and the FKBP12 protein, which is a subunit of the TGF-β1 receptor. These subunits dimerize in the presence of rapamycin with high affinity, yielding a functional SspB adapter that can target tagged proteins for degradation. The system worked perfectly to induce rapid degradation in *E. coli*, and we thought this could be a great tool for dynamic control in *Bacillus subtilis*. A rapid degradation system could be useful for achieving more precise temporal control of perturbations to protein levels, allowing for complex perturbations like oscillations during the bacterial cell cycle, as previously done with inducible promoters^25^. This could enable the investigation of molecular pathways and their dynamics, assess rapid responses to prevent adaptation or masking effects of the stress response, and thereby lead to new mechanistic insights.

We then re-engineered the system to be functional in *Bacillus subtilis*. As depicted in Fig. 2a, the system comprises a split version of SspB fused to the rapamycin-binding domains FKBP12 and FRB. Constitutive promoters of different strengths drive these domains to maintain the proper ratios. Insulators were added to prevent transcriptional readthrough^26^. We shuffled the PV promoters^26^ for ones with different strengths but always preserved the FRB/FKBP12 expression ratio to find the minimum strength that would result in a functional system without overburdening the cell with excess protein. As a comparison, we used the traditional degron system with an inducible promoter expressing SspB. We employed all the different inducible promoters available for B. subtilis with varying levels of leakiness. In addition, we engineered a new, very tight promoter by combining the Phy-spank promoter with the T53C mutation from Pspank*. This promoter has a low dynamic range but is sufficient for degron activation and has zero leakiness. This promoter is referred to here as Phy-spank T35C. This promoter is useful when degrading proteins susceptible to degron leakiness, such as essential genes with low expression. However, it would not have been useful for the CRISPRi system due to its low induction for dCas9.

**Figure 2.**
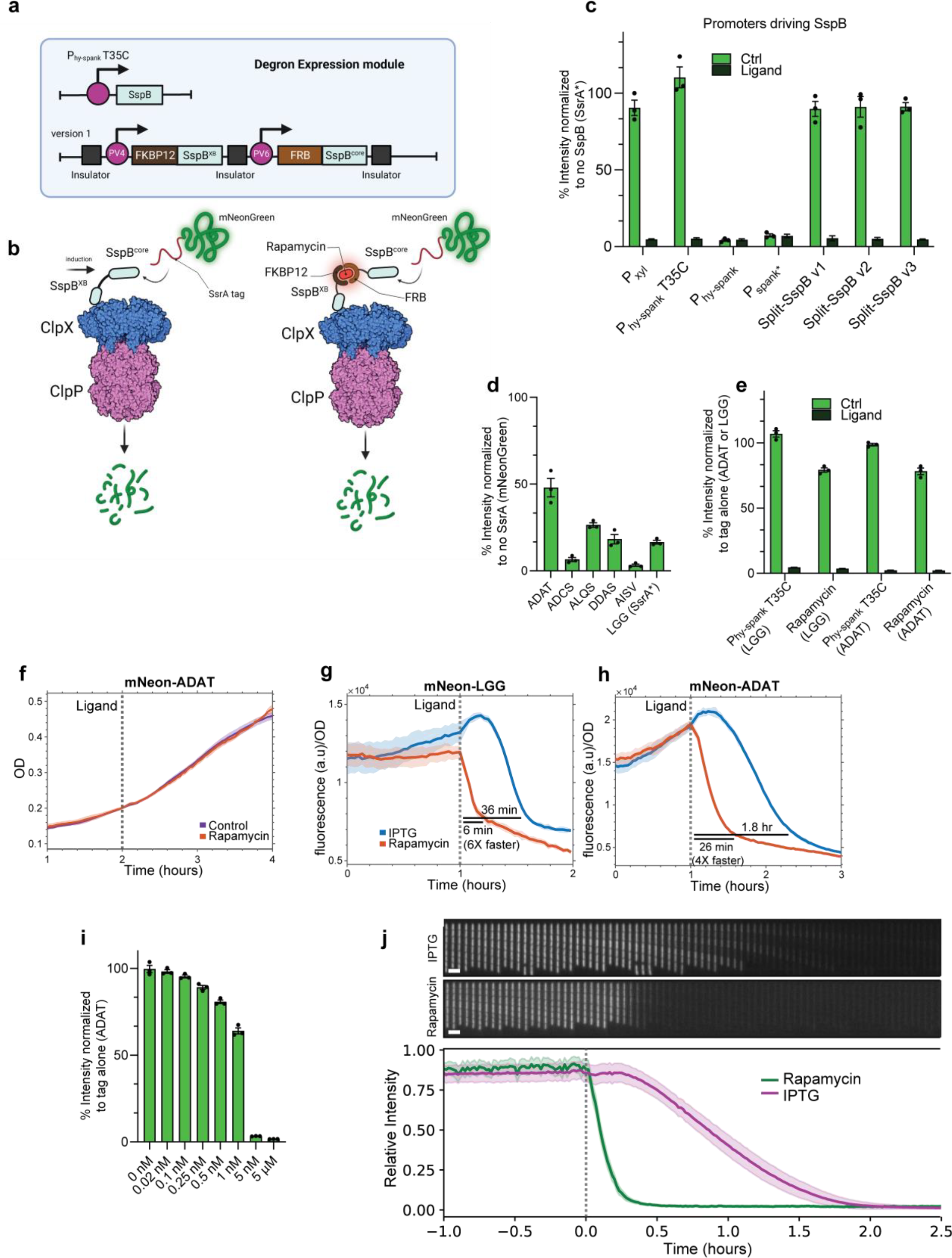
**a**, Schematic diagram of the two degron modalities. The inducible promoter version utilizes Phy-spank T35C to drive full-length SspB expression upon IPTG induction. The rapamycin-induced dimerization construct has been re-engineered for B. subtilis, comprising two constitutive promoters, PV4 and PV6, driving the expression of FKBP12-SspB^XB^ subunit and FRB-SspB^core^ along with genetic insulators. **b**, Diagram representing the molecular mechanism of degradation depending on the construct. In the promoter-inducible version, the protein tagged with SsrA binds to the SspB^core^ subunit and is subsequently carried to ClpX for degradation. In the rapamycin-inducible version, SspB undergoes dimerization in the presence of rapamycin, further binding to the SsrA tag and targeting the protein of interest for degradation. **c**, Flow cytometry quantification of degron fluorescence under different SspB constructs. Ligands (Xylose, IPTG, or Rapamycin) were added accordingly. Rapamycin constructs with different promoter pairs were assessed: v1 = PV4/PV6, v2 = PV5/PV7, v3 = PV3/PV5, where a higher number corresponds to lower expression strength. Results were normalized to mNeon expression without the SsrA tag as a control (n=3 biological replicates). **d**, SsrA tag leakiness screen. Intensities were normalized to mNeon expression without the SsrA tag (n=3 biological replicates). **e**, Comparison of ADAT leakiness and full degradation in Phy-spank T35C and rapamycin constructs versus LGG tag (n=3 biological replicates). **f**, OD600 measurements of control versus rapamycin-treated cells (n=12 technical replicates). **g**, Fluorescence measurements in plate reader format of LGG and **h**, ADAT tagged mNeon green cells. The interval is 2 minutes, and data from the first hour were removed due to low cell density. The dotted line indicates the addition of rapamycin/IPTG. The fluorescence values were normalized by the OD measured simultaneously (n=12 technical replicates). **i**, Fluorescence quantification by flow cytometry of rapamycin titration (n=3 biological replicates). **j**, (Top) Kymographs of a representative Mother Machine lineage of IPTG-inducible degron ADAT and Rapamycin-inducible degron ADAT during the course of a time-lapse experiment. There is a time difference of 3 minutes between each slice of the kymographs. (Bottom) Median of the mean fluorescence intensity per lineage aggregated over all the lineages analyzed under each condition for Rapamycin (green) and IPTG (magenta), normalized with respect to the maximum and minimum mean fluorescence intensities per lineage. The shades represent the mean absolute deviation. At t = 0, we supplemented the media with 5 uM Rapamycin and 100 uM IPTG. The dotted line indicates the time when ligands were added.

Two degradation modes are depicted in Fig. 2b. In the promoter-inducible system, degradation only occurs when the full SspB protein is expressed. As soon as it becomes expressed, it binds to the SsrA tag and brings the protein to ClpX for degradation. On the other hand, the rapamycin system works through dimerization. The SspB subunits tagged with the rapamycin binding domains are constitutively expressed. Once rapamycin enters the system, it serves as a bridge to dimerize SspB subunits, making it functional. SspB can then bring the SsrA tag to ClpX and initiate degradation. The main difference between both systems is that one requires complete transcription and translation of SspB before it can act on the protein. In contrast, the other only requires dimerization, which is a very fast process.

We conducted flow cytometry tests of the different promoters driving SspB and the rapamycin-engineered version with different constitutive promoters of increasing strength (v1, v2, v3). Protein degradation of mNeonGreen as readout occurred in all promoters, successfully eliminating the fluorescent protein (Fig. 2c). For this initial test, we used the SsrA tag LGG, which had been previously chosen as an effective tag in *Bacillus subtilis*^20^. The promoters exhibited different degrees of leakiness when not induced. Of particular interest is the Phy-spank T35C, which had virtually zero leakiness, and was discovered in this study. Phy-spank and Pspank*^20^, thought to have low leakiness, exhibited the highest activity when uninduced, with essentially no protein available in the non-induced state. The Pxyl promoter had relatively similar levels of leakiness to the rapamycin-inducible system. We did not observe any differences in leakiness or degradation when using different constitutive promoters in the rapamycin system. Therefore, we have chosen version 1 as the finalized version, (Fig. 2a).

We then asked whether we could find an alternative SsrA tag with lower leakiness. The LGG tag was initially suggested as the SsrA tag that could provide clear, all-or-none degradation when inducing SspB. However, the previous study that identified LGG as having the least leakiness relied on Western blot experiments^20^. Still, the level of leakiness for LGG is quite significant, making it challenging to tag low-expressed proteins, as it could almost mimic a knockout effect without inducing SspB. We reasoned that by reassessing the tags using a more sensitive technique like flow cytometry, we could quantify the leakiness and potentially observe differences, ultimately identifying a less leaky tag.

We selected the most promising tags proposed in the original study^20^ and compared them to LGG. As a normalizing control, we used the construct without the degradation tag. When measuring the fluorescence with flow cytometry we confirmed that the LGG tag was highly destructive (Fig. 2d). After adding the LGG tag, mNeonGreen levels were reduced to only 20% of the untagged protein levels. For genes with low expression, this could almost eliminate the gene, resulting in lethality or generation of suppressor mutants. This finding may explain why the degron system did not gain popularity in *B. subtilis*. Intriguingly, some of the other tags were equally or even more destructive than LGG, confirming the findings from the previous paper. However, one tag stood out as an exception—the ADAT tag exhibited a basal degradation rate of 50%, preserving almost 30% more protein compared to the LGG tag (Fig. 2d). As a result, we selected the ADAT tag for further testing to assess its potential as a replacement for the standard LGG tag in protein degradation studies.

We then tested whether the ADAT tag was functional in the rapamycin system. When compared to LGG, the ADAT tag performed identically with both the Phy-spank T35C and the rapamycin-inducible system, achieving full degradation from 100% to 0%. Next, we measured degradation dynamics in bulk using plate reader measurements. Since fluorescence increases with increasing cell density, we also measured OD600 for normalization (Fig. f). We observed that the addition of rapamycin did not affect growth. We also found that LGG-tagged mNeonGreen degraded within 5 minutes (Fig. 2g), while ADAT-tagged mNeonGreen degraded within 30 minutes with respect to basal autofluorescence (Fig. 2h). In comparison, IPTG-induced SspB led to tagged mNeonGreen degradation in 30 minutes and 2 hours for the LGG, and ADAT tags, respectively (Fig. 2g, h). This indicates that the ADAT tag leads to slower degradation compared to the LGG tag but it is faster in the rapamycin system compared to the IPTG-induced system. One possible explanation for the slower degradation of the ADAT tag is that it has slower degradation kinetics compared to LGG tag, therefore requiring more time to reach complete degradation. However, it is worth noting that degradation starts almost immediately in the rapamycin system for both tags. In summary, the ADAT tag shares the rapid and complete degradation characteristics of the LGG tag when induced with rapamycin. It proves optimal for degrading genes with low basal expression due to its minimal leaky degradation. Future research could leverage the ADAT tag as a starting point for screening even more efficient tags that maintain higher protein levels with similar response profiles and dynamics.

We conducted a rapamycin titration to determine the minimal dose required for degradation. This information could prove valuable for long live-cell imaging experiments or easier rapamycin wash-out where smaller doses may be necessary. We observed a dose-dependent effect of rapamycin (Fig. 2i) as previously observed in other rapamycin systems^27,28^. This discovery renders the degron system titratable, which adds another advantage to the system. We then investigated the dynamics at the single-cell level. We tested strains with the ADAT tag, and either the rapamycin, or the Phy-spank T35C system in a mother-machine microfluidic device. The mother machine allows for continuous growth, rapid ligand exchange, and the observation of reactions over time^29^, making it ideal for performing complex perturbations like oscillating gene levels^25^. We observed similar temporal responses at the single-cell level, with rapamycin induction resulting in degradation at 30 minutes and IPTG-induced degradation occurring at 2 hours.

Interestingly, we noticed that rapamycin-induced degradation was highly uniform, leading to a sharp decrease in protein levels across all cells in a very fast manner. In contrast, IPTG-induced degradation exhibited more variability, with some cells maintaining mild fluorescence for longer periods, creating greater cell-to-cell variation in degradation. This demonstrates that the rapamycin-inducible system can achieve a precise and rapid knockout of the target gene without residual protein noise. This characteristic could be advantageous for screens, as it provides a clean tool for the targeted destruction of a protein of interest^30^.

## Discussion

In this study, we present a dual system for controlling gene and protein expression in Bacillus subtilis. We have addressed the issue of CRISPRi leakiness, which has long been a challenge in the field. We accomplished this by utilizing a highly inducible and tighter promoter coupled with a degron system that regulates the leakiness of the dCas9 inducible promoter by degrading unwanted dCas9 molecules. We discovered that inducible gRNA allows for the complete elimination of leakiness while also enhancing the CRISPRi effect on highly expressed genes. Additionally, we engineered a fast, low-noise, reversible degradation system through rapamycin-inducible dimerization, enabling rapid resetting of the CRISPRi system without introducing noise. We identified a tight promoter suitable for gene expression studies and a degron tag that preserves protein levels more effectively than previously published tags, particularly for highly sensitive genes. Furthermore, we found that rapamycin titration permits precise control of protein levels through controlled degradation. Collectively, our work represents significant advancements in synthetic biology for CRISPRi and degron systems in Bacillus, opening new avenues for their application in dynamic environments, including microfluidic devices like the mother machine. We anticipate that these engineering tools will facilitate the development of tighter regulatory systems in other organisms, effectively addressing the issue of leakiness and rendering degron systems more versatile. By further optimizing the recently rediscovered tag, it may be possible to minimize degron leakiness, creating a system with zero degradation-related leakiness suitable for genes with very low expression levels. We hope this strategy will enhance and enable better reversible screens in more dynamic environments, ultimately overcoming major limitations associated with these tools.

## Methods

### Culture growth

*B. subtilis* was grown in S750 minimal media supplemented with1% glucose (except for mother machine experiments). Where indicated, xylose, isopropyl thiogalactoside (IPTG) or Rapamycin was added. Cells were grown at 37 °C to an optical density (OD)_600_ of ∼0.4–0.6 unless otherwise noted.

### Strain construction

dCas9 strains were generated upon transformation of PY79 with a 4-5 piece Gibson assembly reaction, amplifying dCas9 from pJMP1 and varying promoters and using mNeonGreen as a fluorescent tag. gRNAs were designed using the software published elsewhere^31^ targeting the mApple gene. The T7 promoter was obtained from the Bacillus Genetic Stock Center item: 1A1613; strain name: LacI-T7-LacZ through an MTA agreement with Rice University. The rapamycin construct was provided as a gift from Joey Davis at MIT. Further optimization of the rapamycin construct to adapt it to *Bacillus subtilis* was done by using genetic parts published previously^26^. When parts were not available from a source, a gBlock was ordered from the sequence. All the genetic designs are available upon request.

### Flow Cytometry

Cells were seeded in 96-well plates, and ligands (xylose, IPTG) were added overnight. On the following day, fresh media was introduced, including ligands if required. Cells were analyzed while in their respective media using an LSRII flow cytometer equipped with laser lines at 488 nm and 561 nm. Post-collection, the cells were subjected to analysis using FlowJo™ v10.9 Software (BD Life Sciences) and gated based on forward scatter/side scatter density in logarithmic scale. The fluorescence levels were quantified as the median of the gated events. Finally, the reported fluorescence values were obtained by subtracting the total cellular fluorescence of a wild-type PY79 strain and normalizing to the control as a percentage.

### Microscopy

For live cell snapshots, cells were grown up to 0.4 OD, pellet and spotted (1 µl) in 1.5% agar pads onto a No. 1.5 glass-bottomed dish (MatTek). Images were collected using a Nikon Eclipse Ti equipped with a Nikon Plan Apo ×60/1.4NA objective and an Andor Clara CCD camera. The light source was ColLED-p300 ultra MB using the 450 nm LED and the 555 nm for green and red fluorescent proteins. We use 200 and 500 msec exposure, respectively.

### Plate reader assays

Cells were seeded in 96-well plates at varying densities and incubated overnight. The following day, the cells were diluted to an optical density of 0.05 at 600 nm (OD600) and subsequently subjected to OD600 measurements and Green fluorescence readings using a Synergy Neo2 plate reader equipped with a filter set configured for Excitation at 479 nm with a 20 nm bandwidth and Emission at 520 nm with a 20 nm bandwidth. Ligands were introduced after 2 hours of growth. To ensure data accuracy, the initial hour of data was omitted due to observed lag time. Data analysis was performed using Matlab.

### Mother machine experiments

MOPS-based minimal media supplemented with 0.2% glucose and 0.05% F108 was used in all stages of this experiment. 5 mL Cultures of bAF739 (Rapamycin-inducible degron), and bAF737 (IPTG-inducible degron) were grown to early stationary phase, then cells were concentrated 10-fold by centrifugation and resuspension in fresh media, and each strain was loaded into separate lanes of a Mother Machine device by centrifugation. Media was then flown through both lanes using different syringe pumps. Once cells grew exponentially (∼3 h after loading) we started imaging them every 1 minute. After 3 hours (at t = 0 in the figure) we switched the media in the lane having bAF739 by media supplemented with 5 uM Rapamycin, and the media in the lane having bAF737 by media supplemented with 100 uM IPTG.

### Imaging analysis

Snapshots were visualized and processed in Fiji. Mother machine image analysis was performed with custom-made software^32^. Briefly, individual lineages were automatically detected and cropped. For each lineage and timepoint, the overall mean fluorescence intensity was computed. This produced a time series of fluorescence intensity per lineage, which was then aggregated over all identified lineages.

## Supplemental tables

Table S1:

*B. subtilis* strains used in this study.

## Author Contributions

A.F.F. designed the experiments, performed genetic cloning, flow cytometry assays, imaging, data analysis and wrote the paper. S.C.H provided intellectual support in genetic design. L.G performed mother machine experiments and analysis and D.E wrote the mother machine analysis code. J.P and E.G provided guidance and helped write the manuscript. All authors approved the final version of the manuscript.

## Notes

### Competing Interest Statement

The authors have declared no competing interest.

## References

1 Hsu, P. D., Lander, E. S. & Zhang, F. Development and applications of CRISPR-Cas9 for genome engineering. Cell 157, 1262–1278 (2014). 10.1016/j.cell.2014.05.010

2 Barrangou, R. & Doudna, J. A. Applications of CRISPR technologies in research and beyond. Nat Biotechnol 34, 933–941 (2016). 10.1038/nbt.3659

3 Sander, J. D. & Joung, J. K. CRISPR-Cas systems for editing, regulating and targeting genomes. Nat Biotechnol 32, 347–355 (2014). 10.1038/nbt.2842

4 Qi, L. S. et al. Repurposing CRISPR as an RNA-guided platform for sequence-specific control of gene expression. Cell 152, 1173–1183 (2013). 10.1016/j.cell.2013.02.022

5 Peters, J. M. et al. A Comprehensive, CRISPR-based Functional Analysis of Essential Genes in Bacteria. Cell 165, 1493–1506 (2016). 10.1016/j.cell.2016.05.003

6 Gilbert, L. A. et al. Genome-Scale CRISPR-Mediated Control of Gene Repression and Activation. Cell 159, 647–661 (2014). 10.1016/j.cell.2014.09.029

7 Li, Y. et al. Metabolic engineering of Escherichia coli using CRISPR-Cas9 meditated genome editing. Metab Eng 31, 13–21 (2015). 10.1016/j.ymben.2015.06.006

8 Utomo, J. C., Hodgins, C. L. & Ro, D. K. Multiplex Genome Editing in Yeast by CRISPR/Cas9 - A Potent and Agile Tool to Reconstruct Complex Metabolic Pathways. Front Plant Sci 12, 719148 (2021). 10.3389/fpls.2021.719148

9 Li, H. et al. Applications of genome editing technology in the targeted therapy of human diseases: mechanisms, advances and prospects. Signal Transduct Target Ther 5, 1 (2020). 10.1038/s41392-019-0089-y

10 Larson, M. H. et al. CRISPR interference (CRISPRi) for sequence-specific control of gene expression. Nat Protoc 8, 2180–2196 (2013). 10.1038/nprot.2013.132

11 Rosano, G. L. & Ceccarelli, E. A. Recombinant protein expression in Escherichia coli: advances and challenges. Front Microbiol 5, 172 (2014). 10.3389/fmicb.2014.00172

12 Martens, K. J. A. et al. Visualisation of dCas9 target search in vivo using an open-microscopy framework. Nat Commun 10, 3552 (2019). 10.1038/s41467-019-11514-0

13 Li, X. T. et al. tCRISPRi: tunable and reversible, one-step control of gene expression. Sci Rep 6, 39076 (2016). 10.1038/srep39076

14 Hawkins, J. S. et al. Mismatch-CRISPRi Reveals the Co-varying Expression-Fitness Relationships of Essential Genes in Escherichia coli and Bacillus subtilis. Cell Syst 11, 523–535 e529 (2020). 10.1016/j.cels.2020.09.009

15 Koopal, B., Kruis, A. J., Claassens, N. J., Nobrega, F. L. & van der Oost, J. Incorporation of a Synthetic Amino Acid into dCas9 Improves Control of Gene Silencing. ACS Synth Biol 8, 216–222 (2019). 10.1021/acssynbio.8b00347

16 Chen, K. N. & Ma, B. G. OptoCRISPRi-HD: Engineering a Bacterial Green-Light-Activated CRISPRi System with a High Dynamic Range. ACS Synth Biol 12, 1708–1715 (2023). 10.1021/acssynbio.3c00035

17 Zetsche, B., Volz, S. E. & Zhang, F. A split-Cas9 architecture for inducible genome editing and transcription modulation. Nat Biotechnol 33, 139–142 (2015). 10.1038/nbt.3149

18 Roth, S., Fulcher, L. J. & Sapkota, G. P. Advances in targeted degradation of endogenous proteins. Cell Mol Life Sci 76, 2761–2777 (2019). 10.1007/s00018-019-03112-6

19 Izert, M. A., Klimecka, M. M. & Gorna, M. W. Applications of Bacterial Degrons and Degraders - Toward Targeted Protein Degradation in Bacteria. Front Mol Biosci 8, 669762 (2021). 10.3389/fmolb.2021.669762

20 Griffith, K. L. & Grossman, A. D. Inducible protein degradation in Bacillus subtilis using heterologous peptide tags and adaptor proteins to target substrates to the protease ClpXP. Mol Microbiol 70, 1012–1025 (2008). 10.1111/j.1365-2958.2008.06467.x

21 Wiktor, J., Lesterlin, C., Sherratt, D. J. & Dekker, C. CRISPR-mediated control of the bacterial initiation of replication. Nucleic Acids Res 44, 3801–3810 (2016). 10.1093/nar/gkw214

22 Davis, J. H., Baker, T. A. & Sauer, R. T. Small-molecule control of protein degradation using split adaptors. ACS Chem Biol 6, 1205–1213 (2011). 10.1021/cb2001389

23 Castillo-Hair, S. M., Fujita, M., Igoshin, O. A. & Tabor, J. J. An Engineered B. subtilis Inducible Promoter System with over 10 000-Fold Dynamic Range. ACS Synth Biol 8, 1673–1678 (2019). 10.1021/acssynbio.8b00469

24 Hambraeus, G., Karhumaa, K. & Rutberg, B. A 5’ stem-loop and ribosome binding but not translation are important for the stability of Bacillus subtilis aprE leader mRNA. Microbiology (Reading) 148, 1795–1803 (2002). 10.1099/00221287-148-6-1795

25 Si, F. et al. Mechanistic Origin of Cell-Size Control and Homeostasis in Bacteria. Curr Biol 29, 1760–1770 e1767 (2019). 10.1016/j.cub.2019.04.062

26 Guiziou, S. et al. A part toolbox to tune genetic expression in Bacillus subtilis. Nucleic Acids Res 44, 7495–7508 (2016). 10.1093/nar/gkw624

27 Mootz, H. D., Blum, E. S., Tyszkiewicz, A. B. & Muir, T. W. Conditional protein splicing: a new tool to control protein structure and function in vitro and in vivo. J Am Chem Soc 125, 10561–10569 (2003). 10.1021/ja0362813

28 Wilmington, S. R. & Matouschek, A. An Inducible System for Rapid Degradation of Specific Cellular Proteins Using Proteasome Adaptors. PLoS One 11, e0152679 (2016). 10.1371/journal.pone.0152679

29 Potvin-Trottier, L., Luro, S. & Paulsson, J. Microfluidics and single-cell microscopy to study stochastic processes in bacteria. Curr Opin Microbiol 43, 186–192 (2018). 10.1016/j.mib.2017.12.004

30 Chassin, H. et al. A modular degron library for synthetic circuits in mammalian cells. Nat Commun 10, 2013 (2019). 10.1038/s41467-019-09974-5

31 Wang, T. et al. Pooled CRISPR interference screening enables genome-scale functional genomics study in bacteria with superior performance. Nat Commun 9, 2475 (2018). 10.1038/s41467-018-04899-x

32 Eaton D. et al. Essentialome-wide Multiplex Imaging Maps Coupling of the Divisome, Cell Envelope and Proteome. (2023 (In preparation)).

